# Sequence variability, constraint and selection in the *CD163* gene in pigs

**DOI:** 10.1101/354159

**Authors:** Martin Johnsson, Roger Ros-Freixedes, Gregor Gorjanc, Matt A. Campbell, Sudhir Naswa, Kimberly Kelly, Jonathon Lightner, Steve Rounsley, John M. Hickey

## Abstract

**Background:** In this paper, we investigate sequence variability, evolutionary constraint, and selection on the *CD163* gene in pigs. The pig *CD163* gene is required for infection by porcine reproductive and respiratory syndrome virus (PRRSV), a serious pathogen with major impact on pig production.

**Results:** We used targeted pooled sequencing of the exons of *CD163* to detect sequence variants in 35,000 pigs of diverse genetic backgrounds and search for potential knock-out variants. We then used whole genome sequence data from three pig lines to calculate a variant intolerance score, which measures the tolerance of genes to protein coding variation, a selection test on protein coding variation over evolutionary time, and haplotype diversity statistics to detect recent selective sweeps during breeding.

**Conclusions:** We performed a deep survey of sequence variation in the *CD163* gene in domestic pigs. We found no potential knock-out variants. *CD163* was moderately intolerant to variation, and showed evidence of positive selection in the lineage leading up to the pig, but no evidence of selective sweeps during breeding.

## Introduction

In this paper, we investigate sequence variability, evolutionary constraint, and selection on the *CD163* gene in pigs. The pig *CD163* gene is required for infection by porcine reproductive and respiratory syndrome virus (PRRSV) [1], a serious pathogen with major impact on pig production [2]. PRRSV-resistant genome-edited pigs with modified *CD163* have been developed, either by knocking out the gene completely or targeting only the fifth scavenger receptor cysteine-rich (SRCR) domain, which is the one that the virus exploits [3–6].

The physiological functions of *CD163* include clearing of haemoglobin from blood plasma [7], adhesion of nucleated red blood cells to macrophages during red blood cell differentiation [8], and immune signalling [9–11]. When red blood cells rupture and haemoglobin is released into the blood stream, haemoglobin is bound by haptoglobin, and the haptoglobin—haemoglobin complex is taken up by macrophages using *CD163* receptors on their surface [7]. Since its natural function is receptor-mediated endocytosis, *CD163* is a target for pathogens entering cells. At least one other virus, the simian hemorragic fever virus [12], has independently evolved to target *CD163*.

Given that genome editing of *CD163* has led to PRRVS-resistant pigs, we wanted to see if natural loss-of-function variants for the *CD163* gene could be identified in elite pigs, in order to investigate the opportunity to develop PPRSV resistance from within existing breeding programs. The aims of this paper were to survey *CD163* sequence variation for such naturally occurring knock-out variants, and to put *CD163* variability in the genomic context of genomic variant intolerance and selection. We used targeted pooled sequencing of the exons of *CD163* to detect sequence variants in 35,000 pigs of diverse genetic backgrounds. We then used whole genome sequence data from three pig lines to put the *CD163* results in the context of the whole genome. We used three complementary population genetic analyses: a variant intolerance score, which measures the tolerance of genes to protein coding variation; a selection test on protein coding variation over evolutionary time; and haplotype diversity statistics to detect recent selective sweeps during breeding.

In summary, our results show that there were no potential knock-out variants in *CD163* in these pigs. Furthermore, *CD163* was moderately intolerant to variation, and showed evidence of positive selection in the lineage leading up to the pig, but no evidence of selective sweeps during breeding.

## Materials and Methods

We used targeted *CD163* exon sequence data from pools of 35,000 pigs of diverse origins, and whole genome sequence data from three lines of pigs from the PIC breeding programme. We used three complementary population genetic analyses: 1) residual variant intolerance score based on segregating exonic SNPs; 2) a gene-based selection test incorporating synonymous and nonsynonymous divergence between the pig and cattle reference genomes; and 3) a selective sweep statistic based on haplotype diversity in imputed whole-genome sequence data from one of the lines.

### Data

We used targeted exon sequencing of the *CD163* gene from 35,000 pigs. The sample included nine lines from the breeding programme of the Pig Improvement Company (PIC). The DNA samples were previously collected in 2011-2016 as part of the operation of the breeding programme.

To put the targeted sequence data in a genomic context, we used whole-genome sequence data from three lines of pigs of the PIC breeding programme. These lines were also sampled in the targeted exome sequencing. We used 1146 individuals from line 1, sequenced at variable coverages. 84 of them were sequenced at 30X coverage, 11 at 10X coverage, 45 at 5X coverage, 561 at 2X coverage, and the remaining 445 at 1X coverage. The individuals and their sequence coverages were chosen with the AlphaSeqOpt algorithm [13,14], with the addition of sires that contributed a large proportion of the genotyped progeny in line 1 that were genotyped as part of the routine breeding activities of PIC. We used 408 individuals from line 2, and 638 individuals from line 3, all of them sequenced at 2X coverage. These individuals were sires that contributed a large proportion of the genotyped progeny in lines 2 and 3 that were genotyped as part of the routine breeding activities of PIC.

### Targeted sequencing of *CD163*

We used a hierarchical pooling strategy to be able to sequence *CD163* exons in many individuals cost-effectively. We pooled 96 DNA samples into one combined DNA sample and constructed a shotgun sequencing library using the ThruPLEX Tag-seq kit from Rubicon Genomics. This kit incorporates unique molecular identifiers that allow for a consensus sequence to be generated from reads originating from the same molecule thus reducing the impact of sequencing errors. 24 such barcoded libraries were combined and used as input into a sequence capture reaction using baits designed against the exons of *CD163* gene (Arbor Biosciences, Ann Arbor, MI). The product of the library capture was then used to generate 2×150bp reads on an Illumina MiSeq sequencer. This pooling scheme allowed us to sequence up to 2304 samples per sequencing run. In total, 35,808 animals were sampled using this scheme.

We aligned reads with bwa mem (v 0.7.15-r1140) [15] against the 10.2 version of the pig genome with an added 33 kbp contig representing the *CD163* genomic region, which was missing from this version of the reference genome. The coding sequence of *CD163* on this contig is identical to the sequence in the version 11.1 of the pig reference genome. We used Connor (https://github.com/umich-brcf-bioinf/Connor) to call consensus sequences from reads with the same unique molecular identifier. We called variants from these consensus alignments using the LoFreq variant caller [16]. We used snpEff [17] to classify the variants as synonymous, nonsynonymous and stop gain variants.

### Validation of potential knock-out variants

The potential stop gain variants detected in the pooled targeted sequencing data were validated by sequencing of individual animals. We went back to the pools where the variants were detected and sequenced amplicons of the appropriate exons from all the samples making up the pool with individual barcodes on the MiSeq. None of the potential stop gain variants were validated by individual sequencing.

### Whole-genome sequence data processing

We aligned reads to the pig genome (Sscrofa11.1) with bwa mem [15], removed duplicates with Picard (https://broadinstitute.github.io/picard/index.html), and called variants with the GATK HaplotypeCaller [18]. We filtered and processed variant call format files with VCFtools [19].

We used the Variant Effect Predictor [20] to find the protein-coding SNPs, and classifying them into synonymous and nonsynonymous SNPs. We used the Ensembl gene annotation [21], version 90. We downloaded variants in *CD163* from the Ensembl variation database.

### Residual variant intolerance score

The residual variant intolerance score [22] measures gene-level tolerance to mutations by counting segregating variants. To calculate the residual variant intolerance, we counted the number of nonsynonymous variants and the total number of variants in each gene, and calculated the studentised residual of the regression between them. We included variants that segregated in at least one of the three lines.

We applied residual variant intolerance score both at the level of the gene, and at the level of the protein domain [23], using protein domains found by identifying Pfam profiles in Ensembl protein sequences with PfamScan [24]. All gene-level analyses were performed on the principal transcript as designated with APPRIS annotation [25].

### Selection analysis in linage leading up to the pig

SnIPRE [26] uses a Poisson model to measure gene-level selection based on between-species divergence and within-species polymorphism. We calculated the divergence between the pig and cattle (UMD 3.1.1) reference genomes using the Nei-Gojobori method [27] which finds the number of potential synonymous and nonsynonymous substitutions between two codons. We aligned the reference genomes using Lastz [28], and refined the alignments using the chain/net method [29]. We excluded all codons that were not fully aligned between genomes, that is, any codon containing an alignment gap or a missing base in any of the genomes. We ran the empirical Bayes implementation of SnIPRE, using the lme4 R package [30].

### Selective sweep analysis by haplotype diversity

We estimated haplotype diversity at CD163, at random 100 control genes of similar length, and at 11 homologs of genes that are stably expressed in humans [31]. The control genes were selected at random from genes of similar genomic length as CD163 (at most 10% difference).

We imputed genome-wide sequence data to 65,000 pigs from line 1, using SNP chip genotypes from 60K or a 15K SNP chip and the line 1 sequence data described above. We extracted mapped read counts supporting each allele from low coverage samples, as outlined in [Ros-Freixedes et al, in prep], and used multilocus hybrid peeling [32], as implemented in AlphaPeel, to phase and impute all individuals to full sequence data in windows around the selected genes.

We extracted all variants falling within exons and introns of the gene in the reference genome and identified haplotypes carried by each individual in each gene, including only genotyped individuals. For variants encompassing each gene, including introns, strings of phased alleles were compared to define haplotypes carried by each individual in each parental chromosome. Strings of alleles that were identical (with a mismatch threshold) between two individuals were considered the same haplotype, and strings with multiple mismatches were considered as different haplotypes. A maximum of two allele mismatches were allowed before two strings were considered different haplotypes to account for sequencing or phasing errors. We then calculated haplotype homozygosities based on the pooled frequency of the two most common haplotypes (H_12_) [33].

### Gene Ontology enrichment

We downloaded Gene Ontology Biological Process terms for Ensembl genes from BioMart [34], and ranked enriched biological processes by the p-value from Fisher’s Exact test.

## Results

### *CD163* sequence variants identified

We used a hierarchical pooling strategy to sequence the exons of *CD163* from over 35,000 pigs from nine lines, and *CD163* variants from whole-genome sequencing of 1146, 638, and 408 pigs from three of the same lines.

Targeted sequencing of exons identified 140 single nucleotide variants in *CD163*. Whole genome sequencing in three lines identified 15 single nucleotide variants in *CD163*, two of which were nonsynonymous, the rest synonymous, and no potential knock-out variants. Table 1 shows the numbers of synonymous and nonsynonymous single nucleotide variants found in each dataset, and the overlap between them. Figure 1 shows the locations of variants in the *CD163* protein sequence and their frequencies in targeted and whole-genome sequencing.

**Table 1:**
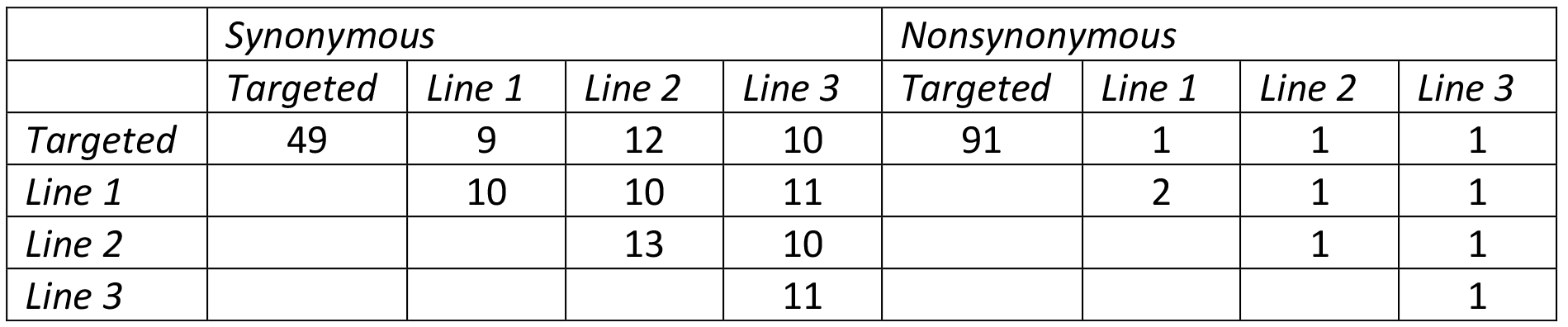
Number of pairwise shared SNPs between variant sets.

**Figure 1:**
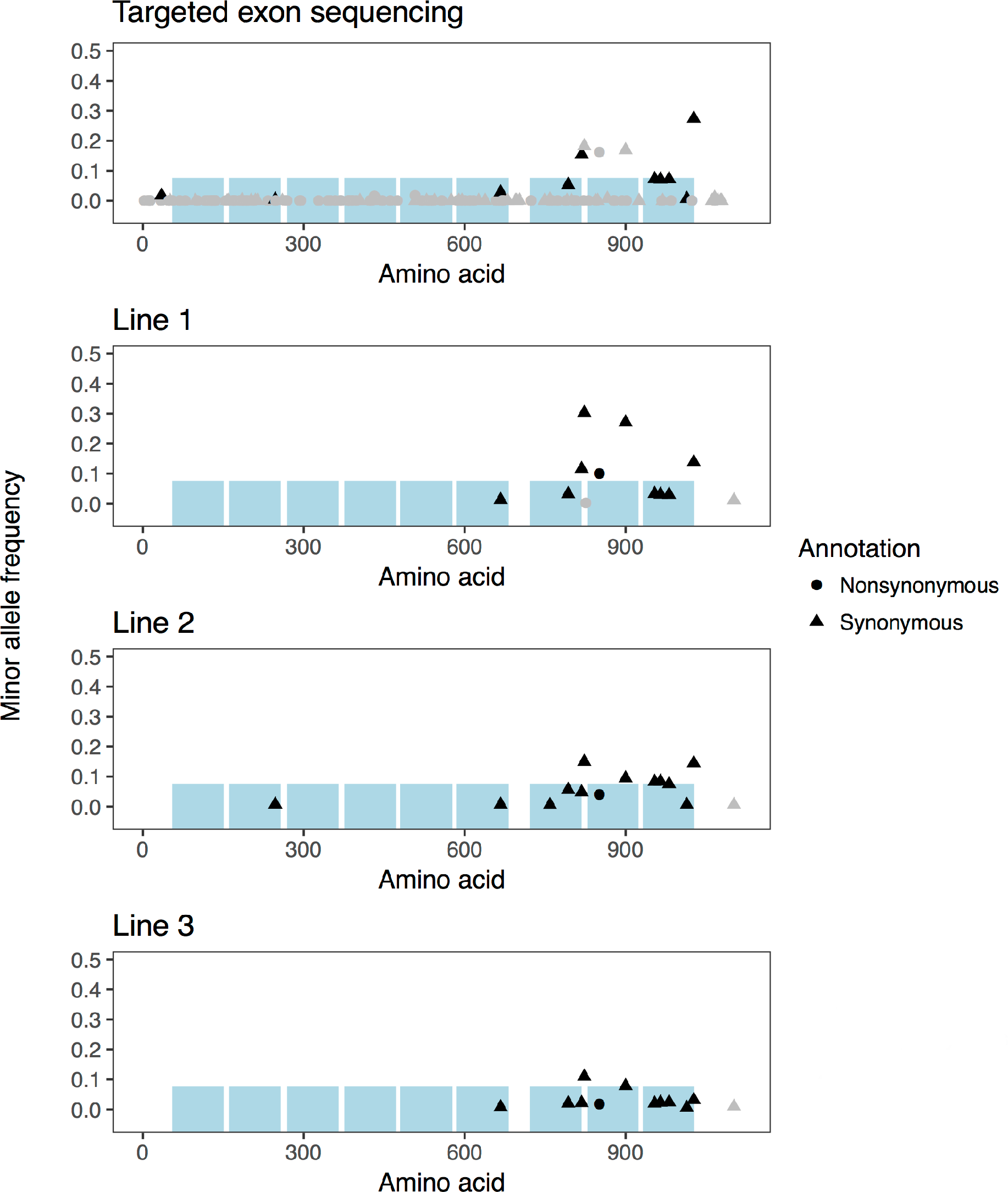
Protein-coding SNPs in CD163 in targeted exon sequencing and whole genome sequencing of three lines. Minor allele frequency showing that discordant variants are rare. Grey and black coloured points indicate replication. Grey dots in targeted sequencing are variants not present in the Ensembl variation database. Grey dots in the whole genome sequenced lines are variants not replicated by the targeted exon sequencing. The blue boxes represent SRCR domains.

The targeted sequencing also identified 14 potential knock-out variants. We followed up on these variants by performing individual sequencing of the animals constituting the pool from which the potential knock-out variant was identified. In this way, we ruled out all the potential knock-out variants as false positives, likely caused by polymerase errors during amplification before incorporation of unique molecular identifiers.

The *CD163* variants detected in whole genome sequence data of the three pig lines were mostly concordant with the targeted sequencing. The discordant SNPs found in the whole genome sequence data were rare. There is one nonsynonymous shared variant, K851R, which occurs at high minor allele frequency. Out of the sequence variants detected, 10 of the variants, most of them at higher frequency, were already present in the Ensembl variation database.

### Residual variant intolerance score

*CD163* was not among the most variant intolerant genes, as measured by a residual variant intolerance score. Figure 2 shows the distributions of gene-level and protein domain-level residual variant intolerance scores with the position of *CD163* and its five variable domains. *CD163* ranked as number 894 out of 17,982 variable autosomal genes. The five variable SRCR domains of *CD163* ranked as 1037 (domain 9), 2686 (domain 7), number 8125 (domains 2 and 6), and 14,147 (domain 8) out of 19,930 variable protein domains, as measured by residual variant intolerance score applied to protein domains identified with the Pfam database.

**Figure 2:**
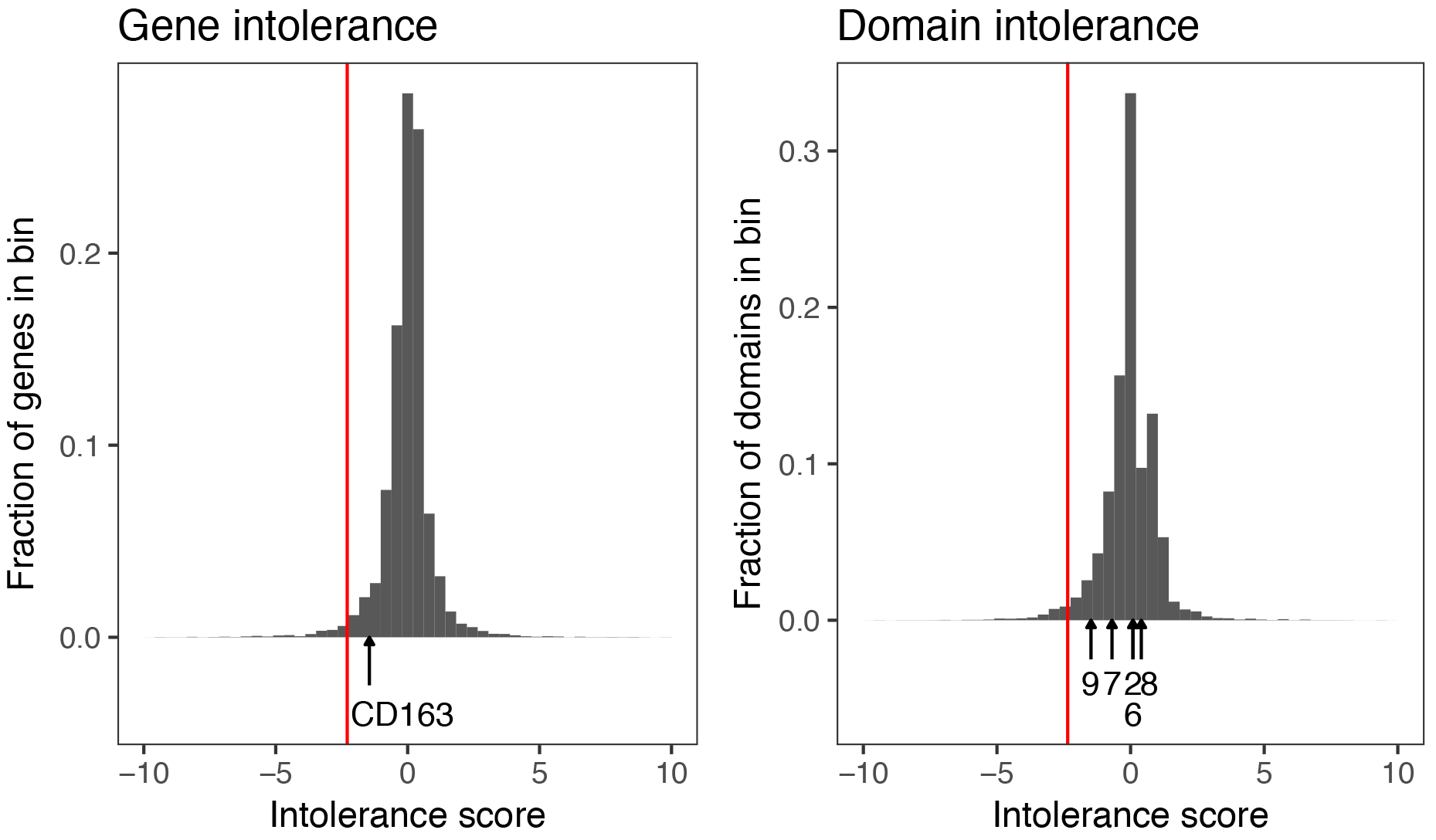
Residual variant intolerance score distributions with CD163 highlighted. The red line is the 2% threshold, and arrows indicate the position of CD163 and five of its SRCR domains.

We used the bottom 2% of the genome-wide residual variant intolerance distribution to highlight 358 variant intolerant genes. They were enriched for basal cellular processes such as microtubule-based movement, cell adhesion, and calcium ion transport (Figure 3).

**Figure 3:**
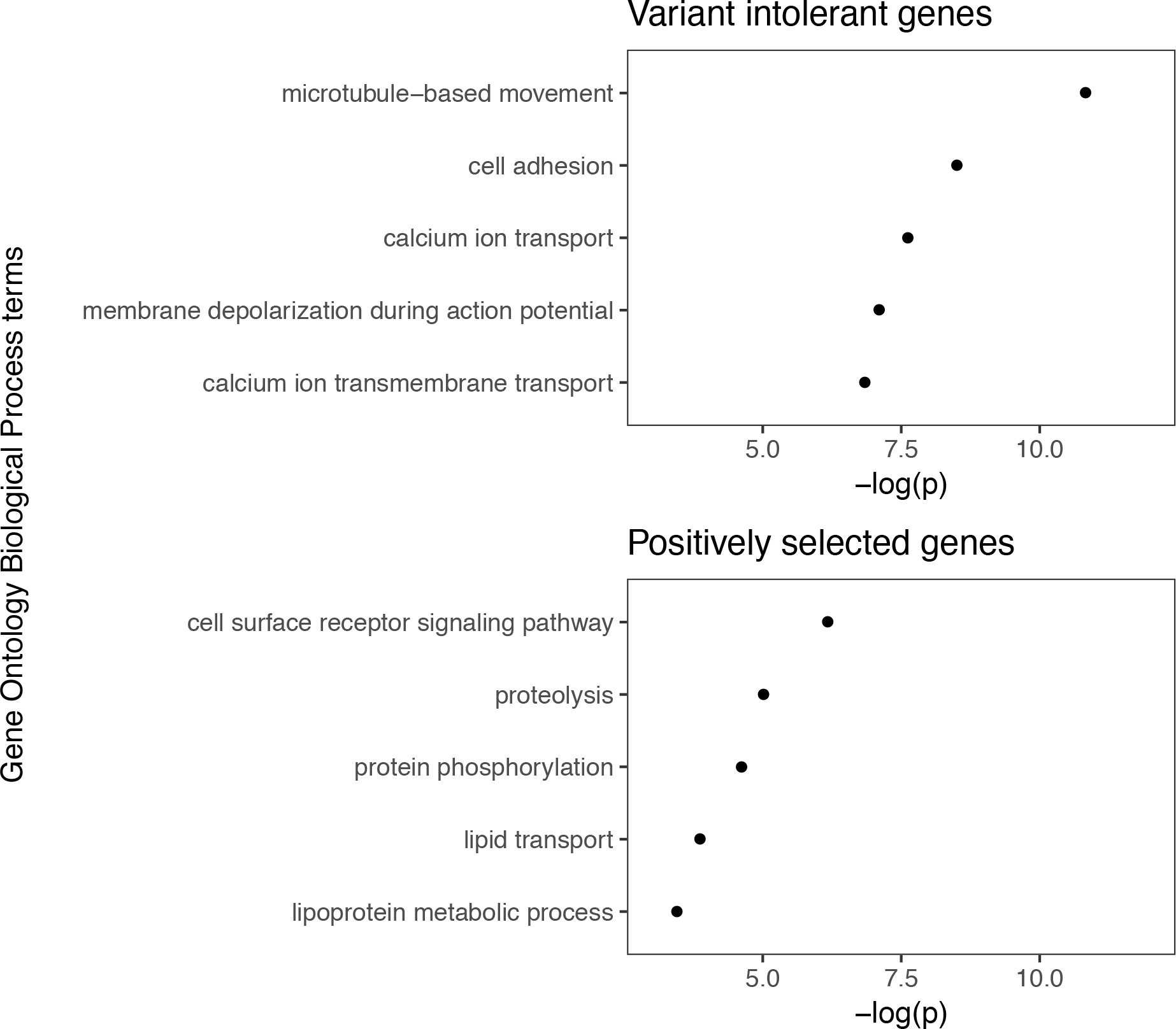
The five most enriched Gene Ontology Biological Process terms in variant intolerant genes andpositively selected genes, with the negative logarithm of the p-value of Fisher’s exact test.

### Selection in the lineage leading up to the pig

*CD163* showed evidence of positive selection in the lineage leading up to the pig, as estimated by the SnIPRE model. Figure 4 shows the selection estimates from the SnIPRE model, highlighting positively and negatively selected genes and the position of *CD163*. We found a total of 1125 putatively selected genes, 778 positively selected genes, and 347 putatively negatively selected. Positively selected genes in the lineage leading up to pig were enriched for cell surface receptor signalling, proteolysis, protein phosphorylation and terms related to lipids (Figure 3).

**Figure 4:**
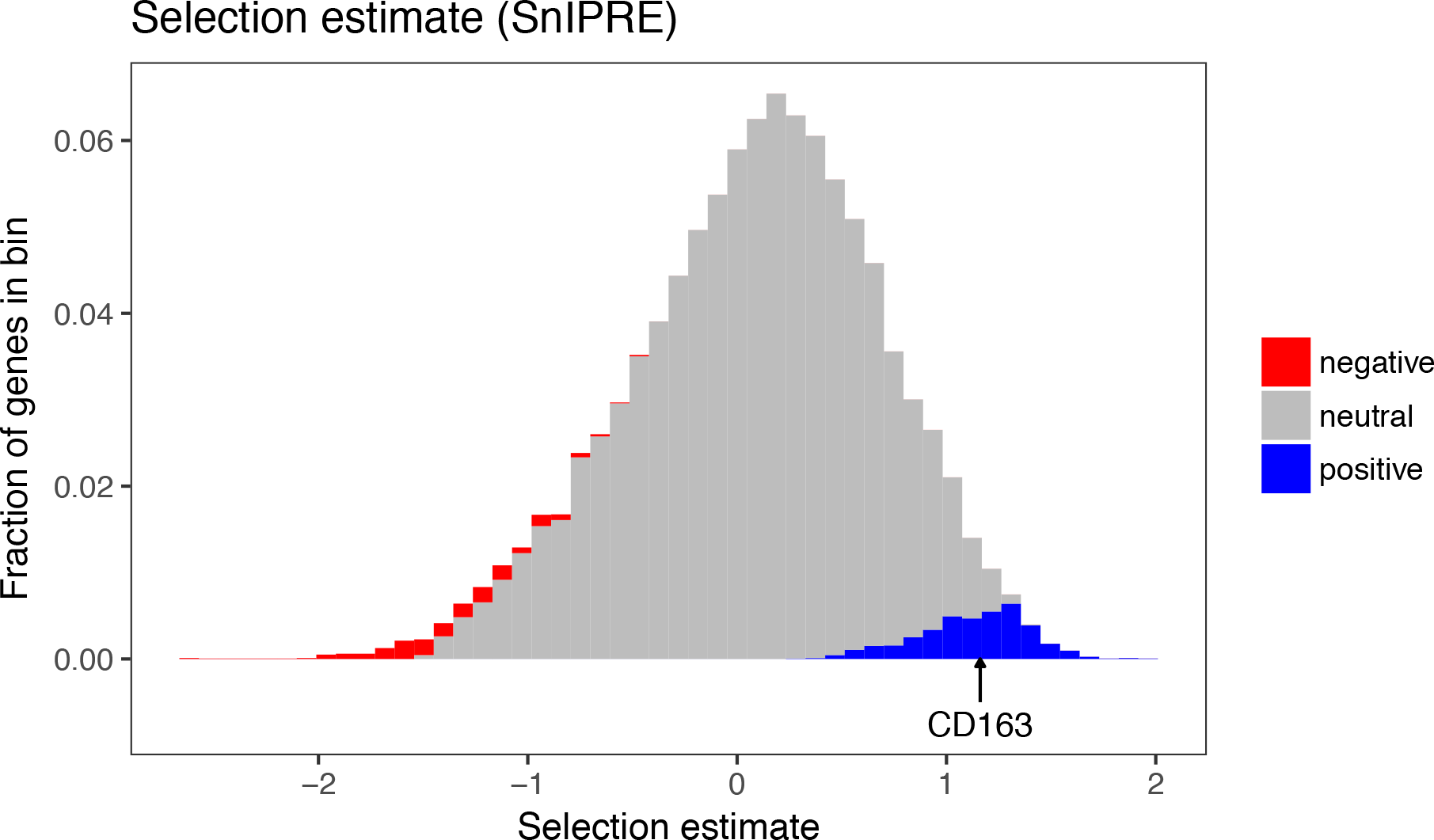
SnIPRE selection estimates. The arrow indicates the position of CD163, which is one of the potentially positively selected (blue) genes.

### Selective sweep analysis

We investigated haplotype diversity in *CD163* in one of the lines using imputed whole genome sequence data. *CD163* showed no evidence of recent selective sweep. We calculated the selective sweep test statistic H_12_, and compared it to 100 randomly selected control genes of similar length. Figure 5 shows H_12_ at *CD163*, the 100 control genes, a set of homologs of genes that are stably expressed in humans, and randomly selected genes labelled as intolerant by their residual variant intolerance score.

**Figure 5:**
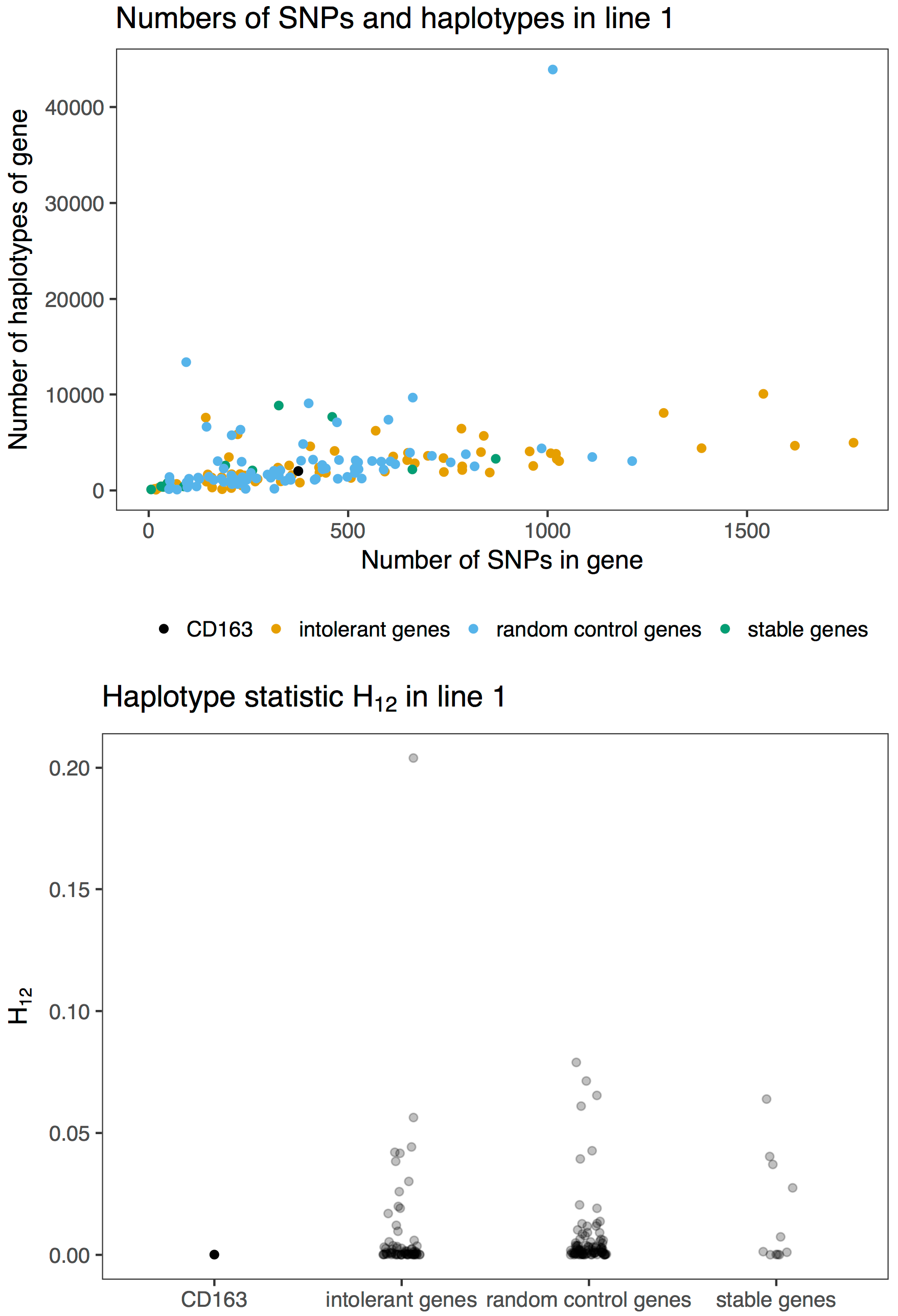
Number of haplotypes, number of SNPs in gene, and selective sweep statistic H_12_ of CD163, 100 control genes of similar length, 100 intolerant genes with low residual variant intolerance score, and 11 control genes that are homologs of human genes with stable expression across many tissues.

## Discussion

In this paper, we investigated sequence variability, evolutionary constraint, and selection on the *CD163* gene in pigs. We identified synonymous and nonsynonymous variants, but no potential knock-out variants in the gene. We found that *CD163* is relatively tolerant to variation, shows evidence of positive selection in the lineage leading up to the pig, and no evidence of selective sweeps during breeding. In the light of these results, we will discuss (i) variant intolerance scores; (ii) selection on *CD163* in the lineage leading up to the pig; (iii) the lack of evidence of selective sweeps; and (iv) technical aspects of the targeted exome sequencing method.

### Residual variant intolerance score

Variant intolerance scores measure the lack or excess of common nonsynonymous variants in a gene [22]. A low variant intolerance score for a gene indicates that its sequence is constrained, and correlates with gene essentiality [35]. The intermediate variant intolerance scores of *CD163* suggest that it is moderately constrained in the pig.

The 2% extreme tail of the variant intolerance distribution was enriched for genes related to microtubule-based movement. This is consistent with an enrichment of microtubule-genes in human essential genes [35].

### Selection in the lineage leading up to the pig

The SnIPRE model is a generalized mixed linear model that estimates the selection effect on each gene with the number of fixed nonsynonymous substitutions compared to an outgroup species [26]. A positive selection estimate means that when comparing the pig to the cow, there has been significantly more nonsynonymous fixed substitutions than expected under neutrality. The synonymous sites are assumed to be under negligible selection. The positive selection effect for *CD163* suggests that its sequence is quite flexible, and has rapidly evolved in the lineage leading up to the pig. Rapid evolution of *CD163* is consistent with its known role in infection. The estimated positive selection on other cell surface genes, including the T cell surface proteins *CD3*, *CD5* and *CD8* and immunoglobulin E receptor *FCER1A*, is consistent with previous observations of rapid immune gene evolution in pigs [36].

### Selective sweep analysis

Selective sweeps occur when the fixation of one or more beneficial variants affects the allele frequencies at linked sites [37]. This signal of recent selection can be detected from population genetics data. When the beneficial variant is already present in the population as standing variation, selection may give rise to a so called soft sweep, which may be more difficult to detect than a sweep arising from a new mutation [38]. Since selection on standing variation is the expectation in animal breeding, we used a statistic designed to detect soft sweeps [33]. The lack of a selective sweep at *CD163* suggest that this gene has not been a target of strong recent selection during pig breeding. However, selective sweep analysis cannot rule the possibility that *CD163* variants could have small effects on some quantitative trait that may have been subjected to subtle allele frequency shifts by selection.

### Technical aspects of the targeted exon sequencing

Targeted exome sequencing of pooled samples is a feasible way to cost efficiently sequence a gene in many individuals. However, as our unsuccessful validation of potential stop gain variants show, this method suffers from low frequency false positives, likely due to polymerase errors before incorporation of unique molecular identifiers. The targeted exon sequencing and whole genome sequencing are in good agreement about higher frequency variants, but since the targeted sequencing sampled a wider span of pig diversity, the rare variant calls may represent genuine rare variants or sequencing errors. However, with the depth of sequencing and validation, we are confident that there are no natural knock-out variants in *CD163* in these pigs.

## Conclusions

We performed a deep survey of sequence variation in the *CD163* gene in domestic pigs. We found no potential knock-out variants. *CD163* was moderately intolerant to variation, and showed evidence of positive selection in the lineage leading up to the pig, but no evidence of selective sweeps during breeding.

## Author’s contributions

SR, MJ, JMH, JL, MAC, and GG conceived the study. SN and KK performed experiments. MJ, RRF, SN and SR analysed data. MJ, JMH, RRF and SR wrote the paper. All authors read and approved the final manuscript.

## Acknowledgements

This work has made use of the resources provided by the Edinburgh Compute and Data Facility (ECDF) (http://www.ecdf.ed.ac.uk).

